# KmerSignificance Score: A discriminative and biologically-informed framework for viral *k*-mer prioritization

**DOI:** 10.64898/2026.05.15.725339

**Authors:** Dylan Lebatteux, Florian Corso, Hugo Soudeyns, Isabelle Boucoiran, Soren Gantt, Abdoulaye Baniré Diallo

## Abstract

Distinguishing closely related viral strains requires identifying genomic regions where subtle sequence differences carry biological significance. While *k*-mer-based approaches offer computational efficiency for genome analysis, existing methods lack standardized frameworks for evaluating which *k*-mers are most informative. Current selection strategies focus primarily on statistical discriminative power without integrating biological relevance. We introduce KmerSignificance Score (KSS), a *k*-mer prioritization framework combining three components: an information-theoretic method measuring strain-distinguishing capacity, an optimized amino acid substitution matrix (MIYATA EVO) for mutation impact assessment, and protein-level functional importance scoring derived from UniProt annotations. KSS produces standardized scores in the [0, 1] interval, enabling direct cross-dataset comparison. The discriminative component achieved classification performance comparable or superior to all tested alternatives (mean F1 = 0.880 vs. 0.718–0.877 for six established methods) while additionally providing bounded scores with consistent empirical distributions for cross-dataset comparability. MIYATA EVO, optimized via genetic algorithm, improved biophysical property correlations by 28.4% over the original MIYATA matrix. Protein scoring on 17,470 viral proteins showed robust agreement with UniProt annotation scores (Kendall *τ* = 0.777) while revealing finer functional distinctions. Literature validation on SARS-CoV-2 (278,738 sequences, 19 variants), HIV-1 (12,223 sequences, 15 subtypes), and human cytomegalovirus (HCMV; 399–646 sequences, 4–8 genotypes) confirmed that high-scoring *k*-mers consistently map to established variant-defining mutations, subtype-specific polymorphisms, and genotype markers. KSS provides a standardized framework for viral *k*-mer prioritization with applications in variant surveillance, molecular epidemiology, and functional annotation. The tool is available at https://github.com/bioinfoUQAM/KmerSignificanceScore.

**Author summary:** Identifying genetic differences between closely related viral strains is essential for pandemic preparedness, vaccine development, and understanding disease outbreaks. With millions of viral genomes now sequenced, researchers need tools that can rapidly pinpoint which genomic differences matter most biologically, not just which are statistically distinctive. Current *k*-mer-based approaches identify patterns that distinguish viral strains but cannot assess whether those differences affect protein function or disease phenotype. We developed KmerSignificance Score (KSS), a framework that we designed to rank short genomic sequences by combining three types of evidence: how well they distinguish viral strains, how much the encoded amino acid changes affect protein properties, and how functionally important the affected protein is. We standardized the resulting scores on a 0-to-1 scale, allowing direct comparison across different viruses and studies. We validated our framework on three major human pathogens (SARS-CoV-2, HIV-1, and human cytomegalovirus) and found that top-scoring positions consistently correspond to sites with documented roles in immune evasion, drug resistance, viral fitness, and strain classification. Our framework can help prioritize genomic features for surveillance of emerging variants, guide experimental validation, and support molecular epidemiology.

## Introduction

High-throughput sequencing technologies have substantially expanded the global virome catalog, with over 6.8 million viral genomes now available in public databases [1]. This extensive diversity presents significant computational challenges for viral characterization and surveillance [2, 3]. Alignment-free approaches using *k*-mers (short nucleotide or amino acid subsequences) have emerged as efficient tools for capturing genetic signatures that distinguish subspecies and strains [4–10].

While *k*-mer-based methods efficiently process large-scale genomic data, determining which *k*-mers merit biological investigation remains challenging. Existing selection strategies suffer from key limitations that reduce interpretability and cross-study applicability. Motif discovery tools like MEME and STREME employ statistical enrichment tests but are designed for binary sequence comparisons rather than multi-class discrimination and often produce redundant motifs [11, 12]. Traditional feature selection frameworks, including univariate filters (chi-squared, mutual information, ANOVA) and machine learning techniques (Random Forest, LASSO), generate dataset-specific score distributions [7, 13–15]. While some metrics like Cramér’s V and normalized mutual information produce theoretically bounded [0, 1] values, their empirical distributions vary substantially across datasets with different class structures, limiting cross-study interpretability. This creates inconsistent *k*-mer rankings and prevents meaningful comparisons between classes or families of viruses.

Most critically, current strategies excel at identifying statistically informative patterns but lack integrated mechanisms to evaluate biological significance. Recent approaches such as GRAMEP [16] illustrate this trend, achieving efficient entropy-based *k*-mer selection for viral variant identification without integrating biological context. This disconnect between computational pattern recognition and functional understanding limits our ability to prioritize *k*-mers associated with mutations affecting pathogen phenotypes such as host adaptation, immune escape, and pathogenicity [17]. Researchers need standardized frameworks that combine discriminative power with biological context to guide experimental validation and functional studies.

To address these limitations, we developed KmerSignificance Score (KSS), a *k*-mer prioritization framework integrating three complementary components.

First, a discriminative component quantifies strain-distinguishing capacity through information-theoretic analysis with adaptive complexity scaling, producing consistent score distributions across diverse classification contexts. Second, an optimized amino acid substitution matrix (MIYATA EVO; see *Materials and methods*), derived from the best-performing matrix among 30 evaluated for biophysical property correlations, improves mutation impact assessment. Third, a protein importance score quantifies functional relevance from UniProt annotations to prioritize variants in well-characterized proteins. KSS generates standardized scores in the [0, 1] interval enabling direct comparison of *k*-mer significance across classes or families of viruses.

We validated each KSS component through comprehensive benchmarking on SARS-CoV-2, HIV-1, and human cytomegalovirus (HCMV), representing complementary evolutionary rates, genome organizations, and classification frameworks. The discriminative component achieved mean classification performance comparable to or above all six established feature selection methods. MIYATA EVO substantially improved biophysical property correlations over the original MIYATA matrix [18]. Protein scoring showed robust agreement with UniProt annotation scores while providing finer granularity. Literature cross-referencing confirmed that top-ranked *k*-mers map to functionally relevant mutations documented in published studies across all three classes of viruses.

## Materials and methods

### KmerSignificance Score framework

KSS evaluates both discriminative capacity and biological significance of *k*-mers through an integrated three-component system. The framework combines: (1) a discriminative component quantifying strain-distinguishing potential through information-theoretic analysis, (2) a mutational component assessing biophysical impact through an optimized amino acid substitution matrix (MIYATA EVO), and (3) a protein component quantifying functional importance from UniProt annotations. Each component generates normalized values in the [0, 1] interval.

Fig 1 illustrates the complete computational workflow comprising five main stages: input processing, sequence alignment and translation, *k*-mer extraction with mutation identification, multi-component scoring, and structured output generation. The following subsections detail each component’s methodology and implementation.

**Fig 1.**
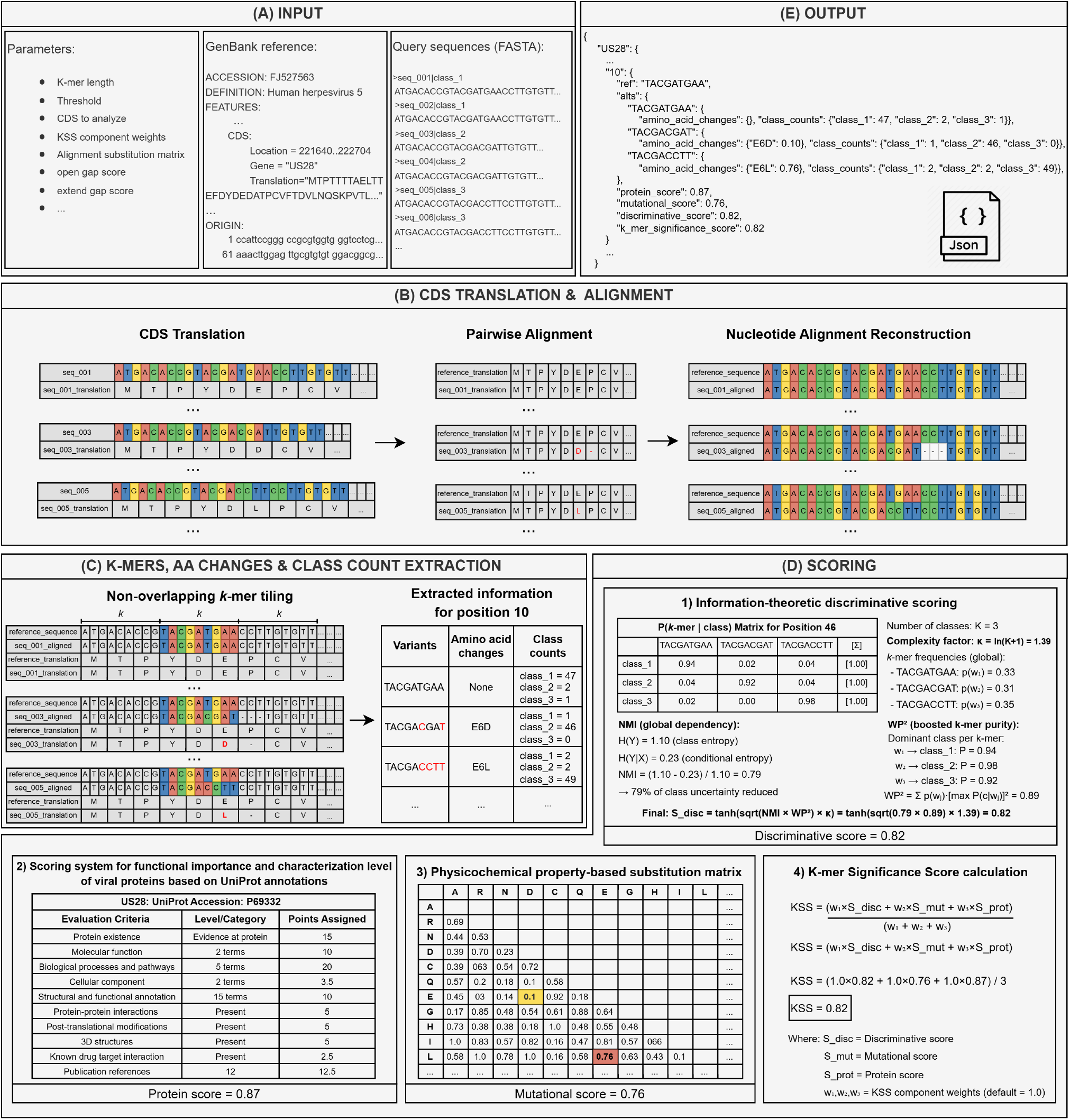
KSS computational workflow. (**A**) Input requirements: user-defined parameters (*k*-mer length, thresholds, component weights), GenBank reference genome with CDS annotations, and query sequences in FASTA format. (**B**) CDS translation and alignment: amino acid sequence generation, pairwise alignment against reference, and nucleotide alignment reconstruction guided by protein alignment. (**C**) *k*-mer extraction using non-overlapping tiling with amino acid change identification and class frequency tabulation for each position. (**D**) Multi-component scoring: (1) information-theoretic discriminative scoring combining normalized mutual information with squared weighted *k*-mer purity, (2) protein functional importance scoring, (3) MIYATA EVO-based mutational impact scoring, and (4) weighted integration into final KSS. (**E**) Structured JSON output containing *k*-mer sequences, component scores, and final KSS values for downstream analysis.

#### Data preprocessing and *k*-mer extraction

KSS requires three inputs: viral coding sequences (CDS) in FASTA format with class labels, a GenBank reference genome, and user-defined parameters. Three key parameters control the analysis: *k*-mer length (*k*), coding sequence selection criteria, and frequency threshold (*t*) representing the minimum proportion of sequences within at least one class that must contain a given *k*-mer. Default parameters were selected based on prior literature and standard alignment practices: nucleotide *k*-mer length *k* = 9, shown to be effective for capturing discriminative viral signatures [6, 7, 19]; frequency threshold *t* = 0.25, ensuring *k*-mers appear in at least 25% of sequences within one viral class (increased to *t* = 0.33 for HIV-1 due to extensive intra-subtype length polymorphisms); and Needleman-Wunsch [20] gap penalties of −10 (open) and −1 (extend). These preprocessing steps establish a consistent coordinate system across viral strains, enabling position-specific calculations.

The preprocessing pipeline comprises four sequential steps:

1. **CDS translation:** Sequences are translated using the standard genetic code to generate amino acid sequences for alignment.
2. **Pairwise alignment:** Amino acid sequences are aligned against GenBank reference translations using the Needleman-Wunsch algorithm [20] with BLOSUM62 matrix, establishing consistent genomic coordinates across strains.
3. **Nucleotide alignment reconstruction:** Guided by amino acid alignments, nucleotide sequences are back-translated by mapping each aligned position to its corresponding codon triplet, maintaining reading frame consistency.
4. **Variant identification and** *k***-mer extraction:** Using adapted KANALYZER functions [21], we extract *k*-mers, variants, amino acid mutations, and class occurrence frequencies from aligned sequences, preparing data for multi-component scoring.

#### Discriminative score

The discriminative component quantifies each position’s ability to distinguish between viral classes through information-theoretic analysis combined with *k*-mer purity assessment. This approach integrates normalized mutual information with class-specific *k*-mer enrichment, scaled by class complexity.

Let 𝒮 = {*s*_1_, …, *s*_*m*_} denote the sequence set and 𝒞 = {*c*_1_, …, *c*_*K*_} the set of *K* viral classes with corresponding labels *y*_*i*_ ∈ 𝒞 for each sequence. For a given position, let *X* ∈ {0, 1}^*m*×*n*^ represent the binary feature matrix where *X*_*ij*_ = 1 if *k*-mer variant *w*_*j*_ is present in sequence *s*_*i*_.

We first compute the entropy of the class distribution:

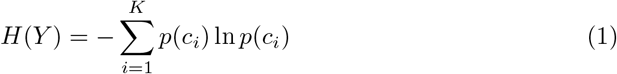

where *p*(*c*_*i*_) = |𝒮_*i*_ |/*m* is the proportion of sequences belonging to class *c*_*i*_.

The conditional entropy *H*(*Y* | *X*) captures the remaining uncertainty about class membership given the *k*-mer configuration:

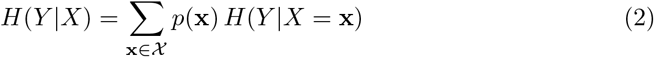

where 𝒳 denotes the set of unique *k*-mer configurations observed across sequences, *p*(**x**) is the proportion of sequences exhibiting configuration **x**, and *H*(*Y* | *X* = **x**) is the class entropy among sequences sharing that configuration.

The normalized mutual information quantifies the proportional reduction in class uncertainty:

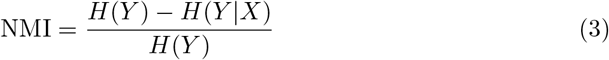

This asymmetric normalization quantifies the fraction of class uncertainty resolved by the *k*-mer configuration, which is appropriate for feature selection where predictive capacity toward the target variable is of primary interest.
To capture class-specific *k*-mer enrichment, we define the purity of each *k*-mer variant as *π*_*j*_ = max_*i*_ *P* (*c*_*i*_ |*w*_*j*_), representing the dominant class proportion among sequences containing variant *w*_*j*_. The squared weighted purity across all variants at the position is then:

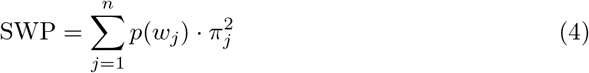

where *p*(*w*_*j*_) is the frequency of *k*-mer variant *w*_*j*_ across all sequences. Squaring the purity term amplifies differences between highly pure *k*-mers (strongly associated with a single class) and moderately pure ones, providing better discrimination between informative and ambiguous positions.

To ensure consistent scoring across datasets with varying numbers of classes, we introduce an adaptive complexity parameter:

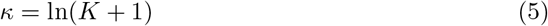

This logarithmic scaling accounts for the inherent difficulty of discrimination tasks: distinguishing among many classes requires stronger signal than binary classification. Table 1 illustrates the complexity values for different class numbers.

**Table 1.** Adaptive complexity parameter *κ* as a function of the number of classes *K*.

| $K$ | $\kappa$ |
| --- | --- |
| 2 | 1.10 |
| 3 | 1.39 |
| 5 | 1.79 |
| 10 | 2.40 |
| 20 | 3.04 |

The final discriminative score combines these components using a geometric mean and applies a hyperbolic tangent transformation to bound the output in [0, 1]:

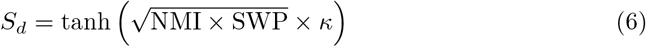

This formulation ensures both information-theoretic and purity components contribute meaningfully through the geometric mean, while the complexity parameter adapts scoring to classification difficulty. The hyperbolic tangent transformation bounds scores in [0, 1] without requiring global dataset normalization. Values approaching 1 indicate positions where *k*-mer patterns strongly predict class membership, while values near 0 correspond to minimal class-distinguishing capacity.

#### Mutational score

Amino acid substitutions influence protein function through alterations in biophysical properties such as size, charge, and polarity [22]. Substitutions between similar residues typically maintain structural stability, while changes involving residues with divergent properties often disrupt function [22–24].

Substitution matrices quantitatively assess these effects by correlating scores with physicochemical properties [23, 25]. While traditional matrices (PAM, BLOSUM) derive from evolutionary frequencies [26, 27], we employ MIYATA EVO, an optimized physicochemical matrix that directly quantifies biophysical dissimilarity. MIYATA EVO was derived by applying genetic algorithm optimization to the original MIYATA matrix [18], maximizing correlation between matrix distances and differences across eight key biophysical properties (described in the following section).

The matrix undergoes two preprocessing steps. First, distance values are inverted so that biochemically similar amino acid pairs receive high values:

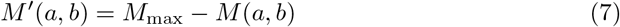

Second, values are normalized to [0, 1] using min-max scaling:

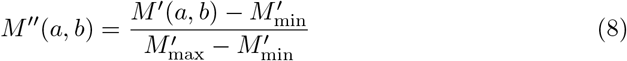

After these transformations, similar amino acid pairs have values approaching 1, while dissimilar pairs approach 0.

For each substitution, the mutational impact is computed as:

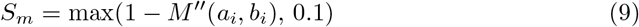

where *a*_*i*_ and *b*_*i*_ represent the original and mutated amino acids. This inversion converts similarity into impact: dissimilar pairs (low *M*″) yield high impact scores, while similar pairs (high *M*″) produce low scores. A minimum threshold of 0.1 ensures that any non-synonymous substitution retains a non-zero contribution to the final KSS. This threshold affects only 8 of 190 amino acid pairs (4.2%), all corresponding to well-established biochemically interchangeable residues (e.g., Leu↔Met, Ile↔Val, Asp↔Glu, Ser↔Thr), and has minimal effect on the final KSS (≤0.033 with equal component weights).

For positions with multiple substitutions, we select the maximum value representing the most biophysically significant change, prioritizing identification of high-impact mutations over averaging that could obscure functionally critical signals. Insertions and deletions receive a configurable score (default 1.0) reflecting their potential for severe structural disruption. This default assumes maximal impact, but users may adjust the indel score to accommodate biological contexts where certain insertions or deletions are less disruptive (e.g., small in-frame indels in flexible loop regions).

#### MIYATA EVO matrix optimization

The mutational component was evaluated by testing 30 substitution matrices from the AAindex database [28] against eight amino acid characteristics. These included seven biophysical properties (Table 2): hydrophobicity, polarity, side chain volume, accessible surface area, isoelectric point, interior-exterior transfer propensity, and aromatic affinity, plus minimum codon distance (MinCodD) as a genetic substitution measure. These characteristics provide comprehensive coverage of physicochemical and genetic factors determining functional consequences of substitutions [29–36].

**Table 2.**
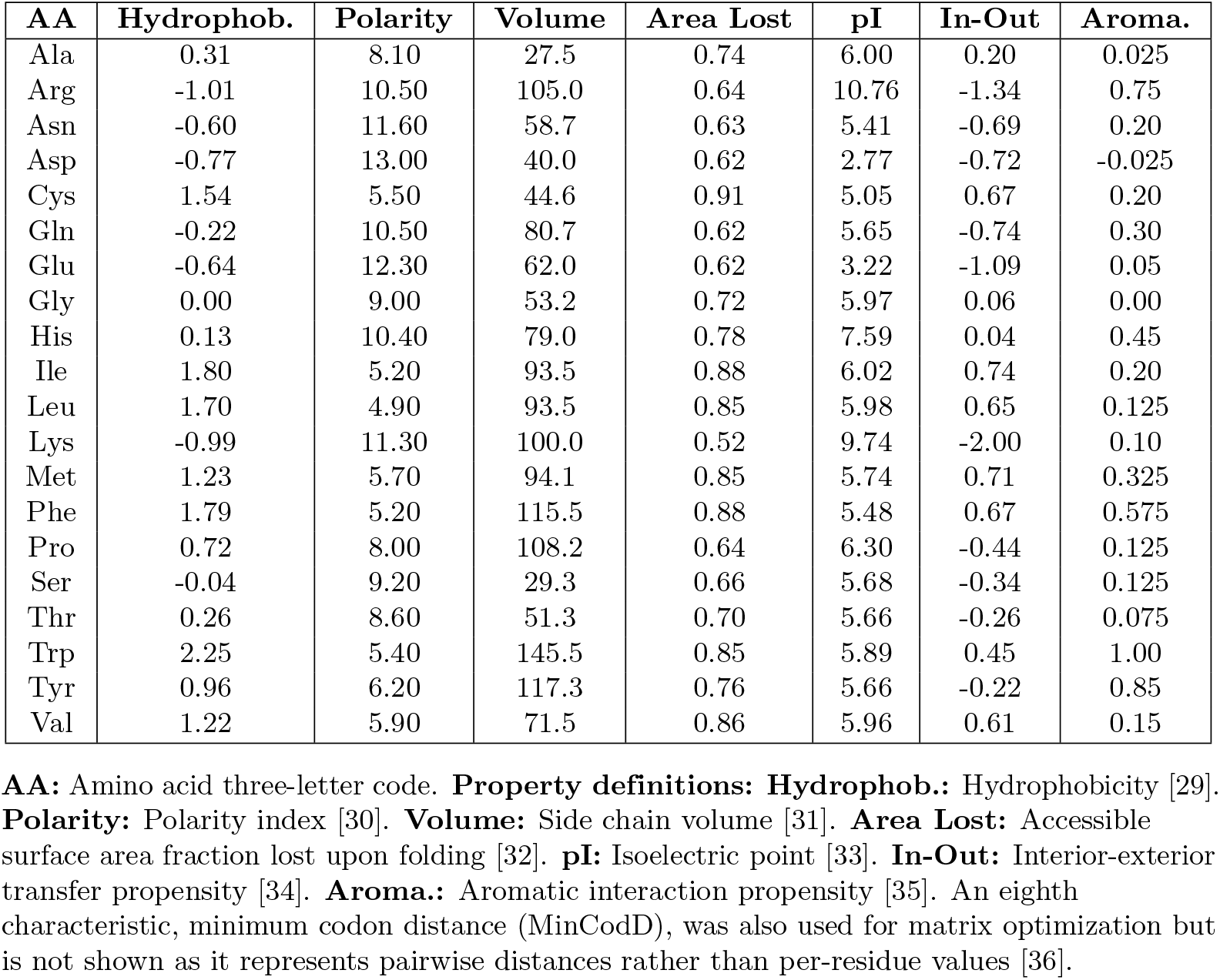
Amino acid biophysical properties for substitution matrix optimization.

For each matrix, substitution values were computed for all 190 unique amino acid pairs (20×19/2). Biophysical distances were calculated as absolute differences between property values for each amino acid pair. Spearman rank correlations between matrix values and biophysical distances were computed for the seven properties, and between matrix values and MinCodD. A composite performance metric summed the eight individual correlations. Statistical significance was assessed using Benjamini-Hochberg false discovery rate (FDR) correction [37] with threshold *q* < 0.01. Results tables report three significance levels: *q* < 0.001 (***), *q* < 0.01 (**), and *q* < 0.05 (*).

The best-performing matrix (MIYATA [18]) was further optimized using a genetic algorithm to maximize the correlation between substitution values and biophysical property differences. The algorithm operated on the 190 unique amino acid pairs (upper triangle), enforcing matrix symmetry by construction and fixing self-substitution distances to zero. Each individual in the population encoded the 190 upper-triangle pairwise distances of the substitution matrix. Offspring were generated through uniform crossover, where each pairwise distance was inherited from one of two parents selected with equal probability (crossover probability 0.5), and fixed-step mutation, where 10% of pairwise distances were randomly perturbed by ±0.1 and clipped to [0, 10]. The algorithm used truncation selection (top 50%) with elitism preserving the best individual. The initial population of 1,000 individuals was generated by mutating the original MIYATA matrix, and the algorithm ran for a maximum of 10,000 generations with early stopping after 100 consecutive generations without improvement (threshold 10^−6^). The fitness function maximized the composite correlation metric. The resulting MIYATA EVO matrix serves as the mutational component, with performance evaluation presented in Results.

#### Protein score

The biological relevance of genetic variants depends on the functional importance of affected proteins [22]. We quantify this through a comprehensive UniProt-based scoring system that prioritizes well-characterized, functionally important proteins.

The framework systematically evaluates ten annotation categories retrieved via UniProt API (S1 Table). These include Gene Ontology terms (molecular function, biological processes, cellular component), protein existence evidence, structural features, protein-protein interactions, post-translational modifications, 3D structures, drug target information, and literature references.

Categories receive differential weights reflecting their predicted relative functional contribution (S1 Table). Gene Ontology terms receive maximum weights as they provide structured, standardized descriptions of protein function [38]. Protein existence evidence and literature references reflect experimental validation confidence and characterization depth, respectively. The remaining categories provide complementary structural, interaction, and pharmacological evidence.

Individual category scores are summed and normalized to [0, 1]:

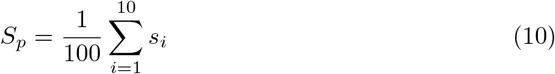

This approach ensures variants in well-characterized, functionally important domains receive higher priority, improving the biological relevance of *k*-mer selection decisions.

#### KSS calculation

The KSS integrates the three components through weighted averaging:

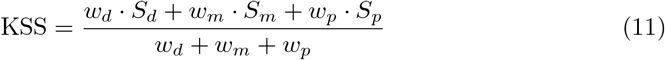

where *S*_*d*_, *S*_*m*_, and *S*_*p*_ represent discriminative, mutational, and protein components, respectively, and *w*_*d*_, *w*_*m*_, *w*_*p*_ are user-defined weights.

Default weights (*w*_*d*_ = *w*_*m*_ = *w*_*p*_ = 1) provide equal contribution from each component. The resulting KSS maintains the [0, 1] interval through component normalization: values approaching 1 indicate *k*-mers with high discriminative capacity and significant mutational impact in well-characterized proteins, while values near 0 correspond to weak discriminative patterns or minimal biological importance.

Weights can be adjusted based on research objectives. Surveillance applications may emphasize discriminative capacity (*w*_*d*_ > *w*_*m*_, *w*_*p*_), while functional studies may prioritize mutational impact (*w*_*m*_ > *w*_*d*_, *w*_*p*_). This flexibility enables tailored prioritization without requiring separate analysis pipelines.

### Validation and benchmarking

#### Viral datasets

We selected three classes of viruses with complementary characteristics for comprehensive KSS validation. SARS-CoV-2 represents rapidly evolving RNA viruses with extensive genomic surveillance data and well-defined variant classifications. HIV-1, a highly diverse retrovirus with complex subtype structure, tests robustness across extreme sequence variation. HCMV represents a DNA virus with moderate evolutionary rates and established genotype-defining mutations enabling targeted validation.

Together, these systems span different evolutionary rates, genome organizations (positive-sense RNA, retrovirus, DNA), and classification frameworks (phylogenetic variants, sequence subtypes, genotype markers). Sequences were collected in November 2025 from GenBank [1] (SARS-CoV-2, HCMV) and the Los Alamos National Laboratory HIV Database [39] (HIV-1).

All available sequences meeting quality control criteria were included without subsampling: (1) exclusion of unverified sequences, (2) sequence length within ±10% of reference, (3) complete coding sequences without premature stop codons, (4) less than 1% ambiguous (non-ACGT) nucleotides, and (5) minimum of 20 sequences per class.

Analysis focused on genes with functional and strain-distinguishing importance: ORF1ab, S, M, N, and E for SARS-CoV-2; gag, pol, and env for HIV-1; UL55, UL73, and US28 for HCMV. Class assignments followed established standards: SARS-CoV-2 variants were classified using Nextclade [40] with Pango lineage nomenclature [41], HIV-1 followed standard subtype definitions [39], and HCMV used genotype classifications [42–44]. Reference genomes for mutation reporting were NC 045512 (SARS-CoV-2), AF033819 (HIV-1), FJ616285 (UL55), and FJ527563 (US28, UL73). Dataset characteristics are summarized in Table 3.

**Table 3.**
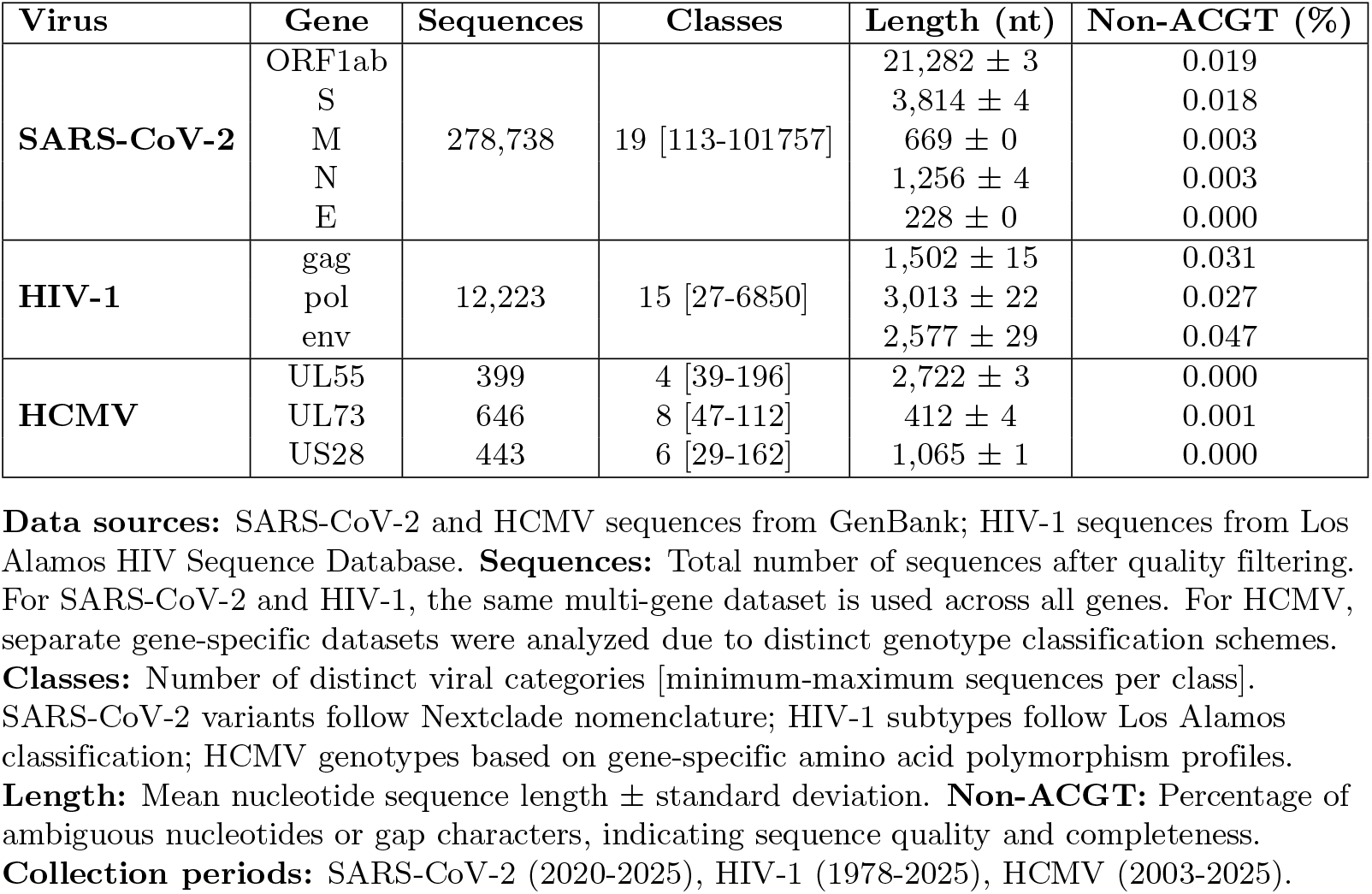
Viral genome datasets used for KSS validation.

| Virus | Gene | Sequences | Classes | Length (nt) | Non-ACGT (%) |
| --- | --- | --- | --- | --- | --- |
| SARS-CoV-2 | ORF1ab | 278,738 | 19 [113-101757] | 21,282 ± 3 | 0.019 |
|  | S |  |  | 3,814 ± 4 | 0.018 |
|  | M |  |  | 669 ± 0 | 0.003 |
|  | N |  |  | 1,256 ± 4 | 0.003 |
|  | E |  |  | 228 ± 0 | 0.000 |
| HIV-1 | gag | 12,223 | 15 [27-6850] | 1,502 ± 15 | 0.031 |
|  | pol |  |  | 3,013 ± 22 | 0.027 |
|  | env |  |  | 2,577 ± 29 | 0.047 |
| HCMV | UL55 | 399 | 4 [39-196] | 2,722 ± 3 | 0.000 |
|  | UL73 | 646 | 8 [47-112] | 412 ± 4 | 0.001 |
|  | US28 | 443 | 6 [29-162] | 1,065 ± 1 | 0.000 |

**Data sources:** SARS-CoV-2 and HCMV sequences from GenBank; HIV-1 sequences from Los Alamos HIV Sequence Database. **Sequences:** Total number of sequences after quality filtering. For SARS-CoV-2 and HIV-1, the same multi-gene dataset is used across all genes. For HCMV, separate gene-specific datasets were analyzed due to distinct genotype classification schemes. **Classes:** Number of distinct viral categories [minimum-maximum sequences per class]. SARS-CoV-2 variants follow Nextclade nomenclature; HIV-1 subtypes follow Los Alamos classification; HCMV genotypes based on gene-specific amino acid polymorphism profiles. **Length:** Mean nucleotide sequence length ± standard deviation. **Non-ACGT:** Percentage of ambiguous nucleotides or gap characters, indicating sequence quality and completeness. **Collection periods:** SARS-CoV-2 (2020-2025), HIV-1 (1978-2025), HCMV (2003-2025).

#### Discriminative component validation

The discriminative component was benchmarked against six established univariate feature selection methods: chi-squared test [45], mutual information (MI) [46], odds ratio [47], Cramér’s V [48], ANOVA F-statistic [49], and normalized mutual information (NMI) [46], along with a random baseline.

Evaluation employed a supervised classification protocol with linear Support Vector Machines (SVM), widely used in *k*-mer-based viral classification studies [6, 7, 50, 51].

For each method, the top-*k* highest-scoring genomic positions were selected, where *k* ∈ {1, 2, 3, 5, 7, 10, 15, 20, 25}. For SARS-CoV-2 and HIV-1, top-*k* positions were selected across all genes simultaneously (pooled analysis), as all genes share the same class structure. For HCMV, top-*k* selection was performed separately for each gene (UL55, UL73, US28) due to distinct genotype classification schemes. Selected positions were used to train classifiers distinguishing viral classes.

To ensure robust estimation, we performed 100 independent iterations with stratified train-test splits. Training set size was determined by adaptive stratified sampling with a target ratio of 70%, constrained between 20 and 500 training sequences per class; remaining sequences were used for testing. Performance was assessed using macro-averaged F1 score, which balances precision and recall across imbalanced class distributions. For each method, mean F1 scores were calculated across all datasets and *k* values to obtain global performance estimates. We also analyzed score distributions across datasets to evaluate whether methods produce consistent, comparable values across classes or families of viruses.

#### Protein score validation

The protein component was evaluated by comparing KSS protein scores with UniProt annotation scores, a heuristic measure combining functional annotation content (e.g., GO terms, functional comments, sequence features) and characterization quality (favoring experimental evidence over predicted annotations). This comparison assessed concordance and identified systematic differences between the two scoring frameworks.

Analysis used 17,470 viral proteins from UniProt (release 2025.06, taxonomy ID 10239) obtained via REST API in JSON format. Spearman rank correlation and Kendall’s tau coefficients evaluated the relationship between UniProt annotation scores and KSS protein scores. Confidence intervals (95%) were calculated using Fisher z-transformation and standard error approximation, respectively. Both methods are appropriate for comparing ordinal with continuous measurement systems.

Granularity analysis assessed KSS discrimination within each UniProt annotation score using descriptive statistics (mean, standard deviation, range, coefficient of variation). Proteins were stratified by annotation scores (1–5) and Kruskal-Wallis tests evaluated whether KSS scores differ significantly across annotation scores. This enables finer prioritization than categorical systems alone, which is particularly useful when selecting proteins for functional studies or identifying well-characterized targets within a given annotation category.

Discordance analysis identified proteins with substantial disagreement between systems. UniProt annotation scores were normalized to [0, 1] using (*s* − 1)/*4* where *s* is the original score. Discordance was quantified as:

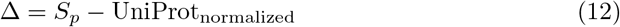

Proteins with |Δ > 0.30| were identified as discordant. This threshold captures moderate-to-large discrepancies while minimizing false positives from minor fluctuations. Cases with Δ >− 0.30 (high KSS relative to UniProt) and Δ *<* − 0.30 (low KSS relative to UniProt) were analyzed to characterize systematic differences between the two frameworks.

#### Functional and discriminative site validation

The integrated KSS framework was evaluated by assessing whether top-ranking positions combine both strain-distinguishing capacity and biological significance. Because KSS integrates discriminative, mutational, and protein components, high-scoring positions should correspond to sites that are simultaneously informative for classification and associated with functionally characterized mutations. To test this, top-ranking genomic positions were compared with experimentally characterized loci across the three classes of viruses.

For SARS-CoV-2 and HIV-1, the top 12 genome-wide positions ranked by KSS were compared against documented functional and strain-defining sites from published literature. To prevent overrepresentation of indel-driven positions, which receive a uniform mutational score of 1.0 and can dominate rankings in highly diverse viruses such as HIV-1, a maximum of three indel positions was permitted within each top-12 selection (≤25% indel representation), ensuring that substitution-based positions, scored by the optimized MIYATA EVO matrix, constitute the majority of each selection. For HCMV, gene-specific analysis identified the top 10 positions per gene (UL55, UL73, US28) with the same indel constraint, due to gene-specific genotyping systems [42–44]. This analysis leveraged substantially larger datasets (399–646 sequences per gene) than previous genotyping studies, enabling evaluation of both KSS performance and established genotyping criteria against expanded sequence diversity.

Literature sources were identified through targeted searches combining viral species, gene names, and specific mutations identified by KSS. Concordance was assessed at exact amino acid positions.

## Results and discussion

### Discriminative component validation

KSS achieved a mean F1 score comparable to the best-performing methods across all datasets and *k* values (0.880), alongside chi-squared (0.877) and odds ratio (0.866), and above NMI (0.861), mutual information (0.857), ANOVA (0.780), Cramér’s V (0.718), and random baseline (0.558) (Fig 2).

**Fig 2.**
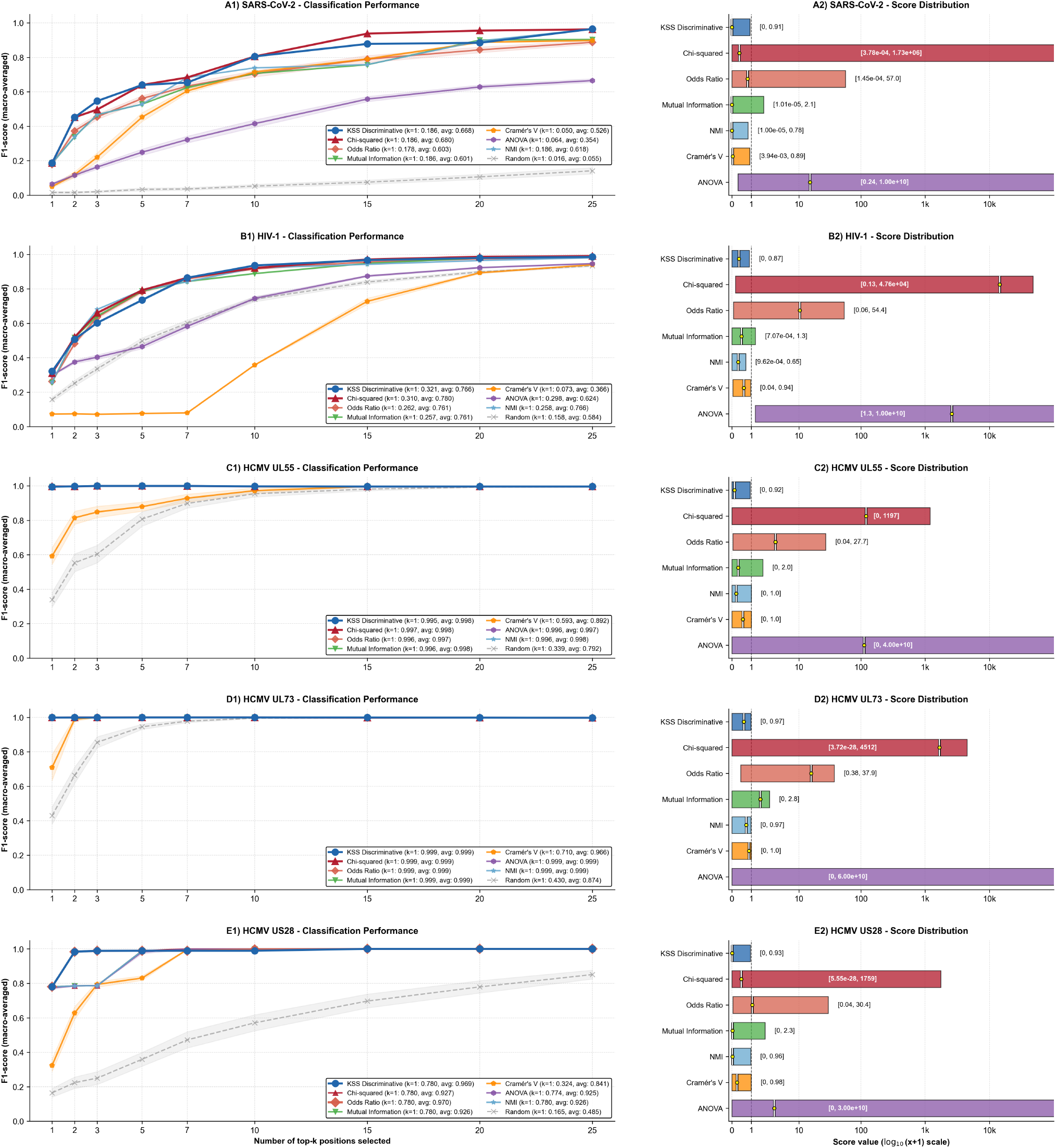
Discriminative component validation across classes and families of viruses. Left panels: Classification performance (macro-averaged F1 score) as a function of the number of top-*k* selected positions for eight feature selection methods. Right panels: Score value distributions showing empirical ranges for each method. KSS discriminative scores (blue) remain bounded within [0, 1] across all datasets, while traditional methods exhibit dataset-dependent scales spanning multiple orders of magnitude. NMI: Normalized Mutual Information. Legend entries show performance at *k* = 1 and mean across all *k* values. Error bands represent 95% confidence intervals from 100 stratified iterations per dataset.

Performance patterns varied between classes and families of viruses reflecting their distinct evolutionary characteristics. For HCMV genes with well-defined genotype structure, all information-theoretic methods achieved near-perfect classification (F1 > 0.99 for UL73, > 0.96 for US28, > 0.92 for UL55), with KSS matching or exceeding alternatives. The clear genotype-defining polymorphisms in these genes provide unambiguous discriminative signals readily captured by multiple approaches.

For HIV-1, characterized by extreme genetic diversity and complex subtype structure, KSS achieved strong performance (mean F1 = 0.766), comparable to chi-squared (0.780) and NMI (0.766), and substantially outperforming Cramér’s V (0.366) and ANOVA (0.624). The moderate differences among top methods likely reflect the inherent difficulty of discriminating 15 subtypes with extensive inter-subtype recombination and intra-subtype diversity.

SARS-CoV-2 presented the most challenging discrimination task despite its lower diversity, due to the 19-class structure with highly imbalanced representation (113 to 101,757 sequences per variant). KSS achieved mean F1 of 0.668, comparable to chi-squared (0.680) and superior to mutual information (0.601), NMI (0.618), and substantially above Cramér’s V (0.526) and ANOVA (0.354). Classification performance improved rapidly with increasing *k*: from F1 = 0.186 with a single position to F1 = 0.965 with 25 positions, demonstrating that discriminative variants are distributed across multiple genomic regions rather than concentrated at single sites.

At *k* = 1, which isolates each method’s ability to identify the single most informative position, KSS achieved the highest F1 for HIV-1 (0.321 vs. 0.310 for chi-squared) and matched the best alternatives for all other datasets. As *k* increases, performance differences between top-ranking methods narrow, consistent with the univariate nature of all evaluated approaches: positions are ranked independently without accounting for information overlap, so methods that identify individually strong but partially redundant positions converge in classification performance.

Beyond classification performance, a critical distinction between KSS and traditional methods lies in score interpretability and cross-dataset comparability. KSS discriminative values remained within the theoretical [0, 1] bounds across all three classes of viruses, with consistent empirical ranges: SARS-CoV-2 [0.00, 0.91], HIV-1 [0.00, 0.87], HCMV UL55 [0.00, 0.92], UL73 [0.00, 0.97], and US28 [0.00, 0.93] (Fig 2, right panels). In contrast, traditional methods produced highly variable, dataset-dependent scales preventing direct cross-study comparison.

Chi-squared values spanned over six orders of magnitude variation between datasets (HCMV UL55: [0, 1,197] vs. SARS-CoV-2: [0, 1.73 × 10^6^]). ANOVA exhibited even more extreme variation, with maximum values reaching 10^10^ across datasets. While theoretically bounded methods (NMI, Cramér’s V) showed more constrained ranges, their empirical distributions varied substantially across datasets. NMI ranged from [0, 0.65] for HIV-1 to [0, 1.0] for HCMV UL55, meaning that a score of 0.5 represents a near-maximum value in one context but a moderate value in another. Cramér’s V showed similarly inconsistent behavior, with mean F1 ranging from 0.366 for HIV-1 to 0.966 for HCMV UL73.

In contrast, KSS consistently utilized the available [0, 1] range across all datasets (empirical maxima between 0.87 and 0.97), ensuring that scores carry comparable meaning regardless of the viral system analyzed. This consistent range utilization enables direct comparison of *k*-mer significance across classes or families of viruses without requiring dataset-specific calibration or normalization, a practical advantage for surveillance applications comparing emerging variants across different pathogens or for meta-analyses integrating results from independent studies.

In summary, the discriminative component achieved competitive classification performance while providing interpretable scores with consistent distributions across diverse classes or families of viruses, addressing a key limitation of existing feature selection approaches for cross-study comparability.

### MIYATA EVO matrix optimization

Among 30 substitution matrices evaluated, the original MIYATA matrix [18] achieved the highest composite value among unmodified matrices (3.566), with seven of eight characteristics showing strong agreement (*q* < 0.001) (Table 4). Hydrophobicity achieved the highest individual correlation (*ρ* = 0.738), while isoelectric point showed non-significant correlation (*ρ* = 0.130, *q >* 0.05), indicating optimization potential.

**Table 4.** MIYATA EVO matrix optimization and validation. Comparison of original MIYATA with genetically optimized MIYATA EVO and top-ranked substitution matrices from AAindex database.

| Rank | Matrix | Hydro. | Polar. | Volume | Area | pI | In-out | Aroma. | MinCodD | Composite |
| --- | --- | --- | --- | --- | --- | --- | --- | --- | --- | --- |
| 1 | MIYATA_EVO | 0.849*** | 0.791*** | 0.286*** | 0.764*** | 0.419*** | 0.773*** | 0.284*** | 0.412*** | **4.578** |
| 2 | MIYATA | 0.738*** | 0.727*** | 0.398*** | 0.554*** | 0.130 | 0.471*** | 0.277*** | 0.270*** | 3.566 |
| 3 | FENG | 0.617*** | 0.628*** | 0.216** | 0.524*** | 0.163* | 0.416*** | 0.253*** | 0.707*** | 3.525 |
| 4 | BENNER74 | 0.697*** | 0.666*** | 0.302*** | 0.621*** | 0.006 | 0.433*** | 0.289*** | 0.427*** | 3.441 |
| 5 | BENNER22 | 0.674*** | 0.638*** | 0.283*** | 0.550*** | 0.031 | 0.397*** | 0.321*** | 0.544*** | 3.439 |
| 6 | GONNET1992 | 0.705*** | 0.670*** | 0.295*** | 0.629*** | −0.001 | 0.440*** | 0.276*** | 0.388*** | 3.402 |
| 7 | VTML250 | 0.677*** | 0.638*** | 0.311*** | 0.595*** | 0.021 | 0.438*** | 0.302*** | 0.370*** | 3.351 |
| 8 | JONES | 0.626*** | 0.600*** | 0.245*** | 0.503*** | 0.043 | 0.389*** | 0.303*** | 0.601*** | 3.309 |
| 9 | BLOSUM62 | 0.678*** | 0.660*** | 0.247*** | 0.592*** | 0.090 | 0.425*** | 0.240*** | 0.339*** | 3.271 |
| 10 | VTML160 | 0.638*** | 0.603*** | 0.306*** | 0.560*** | 0.030 | 0.412*** | 0.318*** | 0.392*** | 3.257 |

**Hydro.:** Hydrophobicity. **Polar.:** Polarity. **Area:** Fraction of accessible surface area lost upon folding. **pI:** Isoelectric point. **In-out:** Interior-exterior propensity. **Aroma.:** Aromaphilicity. **MinCodD:** Minimum codon distance. **Composite:** Sum of all eight correlation coefficients. All values represent Spearman rank correlation coefficients ( $\rho$ ) calculated across 190 amino acid pairs. \*\*\* $q < 0.001$ , \*\* $q < 0.01$ , \* $q < 0.05$ after Benjamini-Hochberg FDR correction. MIYATA\_EVO was generated via genetic algorithm optimization (population = 1,000, max generations = 10,000, mutation rate = 0.1, crossover probability = 0.5) operating on 190 unique pairs (upper triangle) to maximize composite score while enforcing symmetry by construction. Early stopping after 625 generations following 100 consecutive generations without improvement. Improvement over MIYATA: +28.4%.

Genetic algorithm optimization of the MIYATA matrix (see Materials and methods) converged after 625 generations, yielding MIYATA EVO with composite value 4.578, a 28.4% improvement over the original.

MIYATA EVO demonstrated substantial improvements across seven of eight characteristics (Table 4). All eight relationships achieved the highest significance level (*q* < 0.001), compared to only seven for the original. The largest relative gains occurred for isoelectric point (+222.3%), interior-exterior propensity (+64.1%), MinCodD (+52.6%), accessible surface area (+37.9%), and hydrophobicity (+15.0%). Polarity (+8.8%) and aromaphilicity (+2.5%) showed moderate improvements. Side chain volume decreased (*ρ* = 0.286 vs. 0.398, −28.1%) but remained highly significant (*q* < 0.001), indicating the algorithm prioritized overall composite performance rather than individual maxima. MIYATA EVO exceeded the second-best matrix (MIYATA original) by 28.4% and the third-best (FENG) [52] by 29.9%. Notably, several top-ranked matrices showed non-significant isoelectric point values after FDR correction, whereas MIYATA EVO maintained *q* < 0.001 across all characteristics.

The improved biophysical correlations of MIYATA EVO have practical implications for viral mutation assessment. BLOSUM and PAM matrices derive substitution scores from observed evolutionary frequencies across diverse protein families over millions of years [26, 27]. These scores reflect how often substitutions occur during gradual sequence divergence, favoring conservative replacements that maintain function. Viral proteins, however, evolve under distinct selective pressures operating on shorter timescales. Immune selection, drug resistance, and host adaptation drive evolution over months to years [17, 53], where substitutions are selected for immediate functional consequences rather than long-term evolutionary compatibility. MIYATA EVO’s physicochemical foundation, which directly quantifies biophysical property differences, may better capture functionally consequential viral mutations than frequency-based matrices. This distinction may be particularly relevant for surveillance: rare substitutions causing substantial charge or structural alterations may be strongly selected for immune escape despite their evolutionary rarity, and MIYATA EVO can flag such high-impact mutations based on biochemical properties rather than historical frequency patterns. Specifically, MIYATA EVO’s improved correlations for isoelectric point, interior-exterior propensity, and accessible surface area enhance assessment of substitutions affecting charge distribution, membrane interactions, and conformational stability. These properties are particularly relevant for viral envelope and surface proteins where mutations modulate immune recognition while maintaining structural integrity [22]. The decrease in side chain volume correlation (*ρ* = 0.286 vs. 0.398, −28.1%), which remained highly significant (*q* < 0.001), represents a trade-off as the algorithm prioritized gains in properties with greater relevance to viral protein function.

### Protein score validation

Across 17,470 viral proteins, KSS values ranged from 0.000 to 1.000 (mean = 0.313, SD = 0.217, median = 0.300), demonstrating full utilization of the available range. Proteins distributed across UniProt’s five annotation scores [54]: score 1 (*n* = 4, 387, 25.1%), score 2 (*n* = 4, 937, 28.3%), score 3 (*n* = 4, 121, 23.6%), score 4 (*n* = 2, 140, 12.2%), and score 5 (*n* = 1, 885, 10.8%).

KSS showed strong concordance with UniProt classifications. Spearman rank correlation revealed strong positive association (*ρ* = 0.900, 95% CI: 0.897-0.903, *p* < 0.001). Kendall’s tau yielded *τ* = 0.777 (95% CI: 0.767-0.786, *p* < 0.001), confirming robust agreement while accounting for tied ranks. Mean KSS values increased monotonically across scores, from 0.066 in score 1 to 0.668 in score 5. No confidence intervals overlapped between adjacent scores (Table 5). Kruskal-Wallis test confirmed highly significant differences (*H* = 14,197.59, df = 4, *p* < 0.001).

**Table 5.**
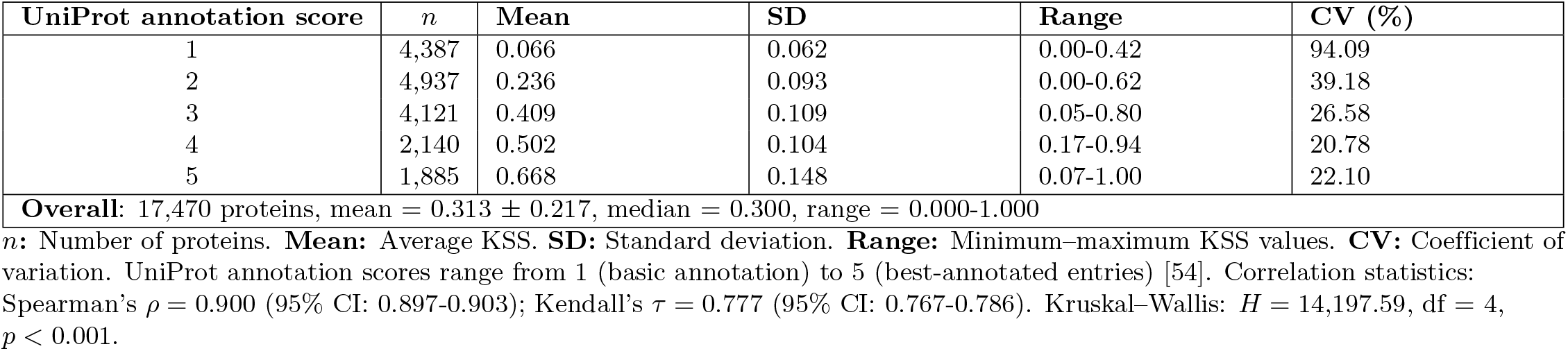
Protein component validation across UniProt annotation scores. KSS distribution statistics for 17,470 viral proteins stratified by UniProt annotation scores.

Coefficient of variation (CV) analysis revealed differential variability across scores. Score 1 exhibited the highest CV (94.09%), largely a mathematical consequence of its near-zero mean (0.066): even modest absolute dispersion (SD = 0.062) produces high relative variation when the denominator approaches zero. Biologically, this reflects the heterogeneity within score 1, which groups completely unannotated proteins (KSS = 0.00) alongside proteins with partial experimental evidence (up to KSS = 0.42), differences that KSS captures but UniProt’s categorical system does not. CV decreased progressively through scores 2 (39.18%) and 3 (26.58%), reaching minimum at score 4 (20.78%), indicating greater scoring consistency for well-characterized proteins. Score 5 showed slight CV increase to 22.10%, reflecting variability in functional complexity even among extensively annotated proteins.

Discordance analysis identified proteins with substantial disagreement between systems. Using threshold |Δ| > 0.30 (where Δ = KSS minus normalized UniProt value), 23 proteins (0.1%) exhibited high positive discordance (high KSS relative to UniProt). In contrast, 1,783 proteins (10.2%) showed high negative discordance (low KSS relative to UniProt). This approximately 78:1 asymmetry reflects systematic differences in scoring philosophy.

While UniProt employs qualitative curation standards emphasizing completeness and evidence quality [55], KSS translates similar principles into a quantitative scoring framework, weighting functional annotations and experimental evidence most heavily.

Analysis of extreme discordance cases (S2 and S3 Tables) reveals that positive discordance proteins (high KSS, low UniProt score) consistently show experimental validation at protein level (protein existence PE=1 in all cases) with experimentally determined 3D structures (70%), despite minimal GO annotations. In contrast, negative discordance proteins (high UniProt score, low KSS) nearly all lack experimental structures (5%) and are predominantly inferred by homology (PE=3 in 70%). For instance, the most extreme case of positive discordance is HCMV protein UL78 (P16751, Δ = +0.42), which possesses experimental evidence, seven structural/functional features, characterized post-translational modifications, and a cryo-EM structure despite minimal UniProt annotation, with recent discoveries validating its G protein-coupled receptor (GPCR) activity [56, 57]. Conversely, the most extreme case of negative discordance is Sulfolobus islandicus rod-shaped virus 1 protein 1070 (Q8QL26, Δ = −0.93), which received maximum UniProt score yet minimal KSS due to predicted status (PE=4), absent structure, and zero functional annotations. These patterns confirm KSS prioritizes experimental validation, functional characterization depth, and structural evidence over annotation completeness.

The strong relationships (*ρ* = 0.900, *τ* = 0.777) and monotonic progression demonstrate that KSS captures annotation quality patterns consistent with UniProt’s categorical system while providing continuous, granular values enabling finer prioritization. The discordance patterns reveal complementary information. UniProt emphasizes annotation completeness while KSS emphasizes functional characterization depth and experimental validation. This complementarity suggests KSS can identify functionally important but under-annotated proteins for prioritization, as well as extensively annotated proteins that may benefit from additional experimental characterization.

### Functional and discriminative site validation

The integrated KSS framework was validated by comparing top-ranking positions with established functional and strain-distinguishing loci across all three classes of viruses (Fig 3).

**Fig 3.**
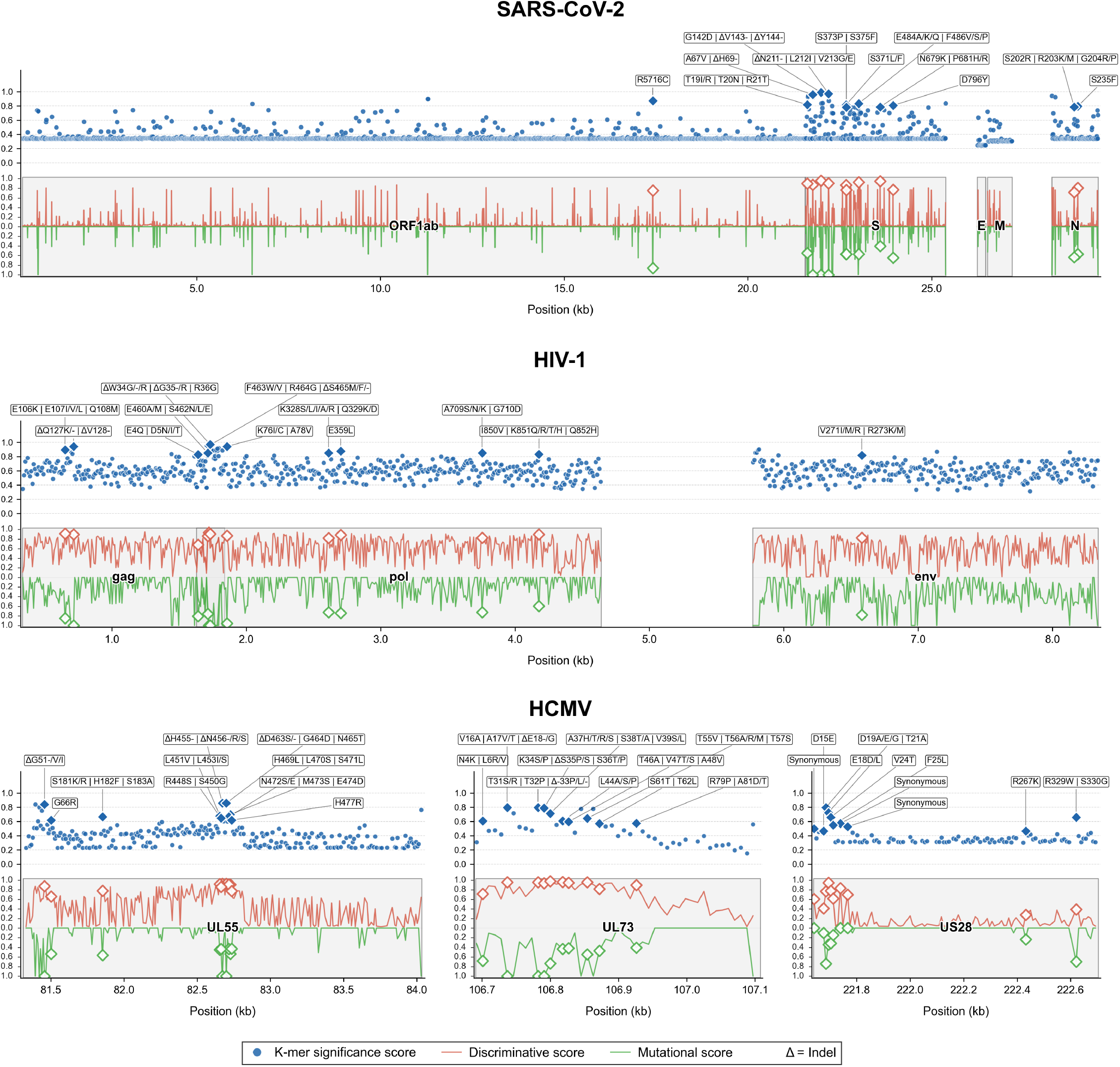
Functional and discriminative site validation across classes and families of viruses. KSS (blue), discriminative component (red), and mutational component (green) across genomic coordinates for SARS-CoV-2, HIV-1, and HCMV. Gene annotations shown as gray boxes. Top-ranking sites are marked with diamond symbols on each score component (blue: KSS, red: discriminative, green: mutational). Top twelve positions for SARS-CoV-2 and HIV-1 (genome-wide) and top ten per gene for HCMV, selected with a maximum of three indel positions per selection. See S4–S6 Tables for detailed mutation listings and literature references.

For SARS-CoV-2, the twelve top-ranking positions (KSS = 0.78–0.99) distributed across three genes: nine in Spike (S), two in Nucleocapsid (N), and one in ORF1ab. The three highest-scoring positions, all involving indels (mutational score = 1.0), localized to the Spike N-terminal domain (NTD). Positions 142–144 (KSS = 0.99; G142D, V143del, Y144del) and 211–213 (KSS = 0.97; N211del, L212I, V213G) correspond to primary deletion clusters defining Omicron, Delta, and Alpha lineages within the NTD antigenic supersite targeted by neutralizing antibodies [58–60]. Positions 67–69 (KSS = 0.96; H69del, A67V) correspond to recurrent deletion region 1 (RDR1), associated with immune escape and antibody evasion [58, 61].

The nine substitution-based positions extended across six functionally distinct genomic regions. Three positions clustered within the receptor-binding domain (RBD): E484A/F486V (KSS = 0.83) encompasses the extensively characterized E484A/K immune escape mutation reducing neutralization by convalescent and vaccine sera [59]; S371L/F (KSS = 0.81) alters inter-protomer RBD interactions and spike conformational dynamics, defining all Omicron BA.2+ sublineages; and S373P/S375F (KSS = 0.78) completes the 371–375 mutation cluster characteristic of Omicron [61]. At the furin cleavage site, P681H/R (KSS = 0.79) enhances Spike processing and defines Alpha (P681H), Delta (P681R), and subsequent lineages [61]. In the NTD, T19I (KSS = 0.82) contributes to antigenic remodeling in BA.2+ sublineages. In the S2 subunit, D796Y (KSS = 0.81) marks virtually all post-ancestral Omicron lineages. In the Nucleocapsid, R203K/G204R (KSS = 0.79) in the serine/arginine-rich linker represents variant-defining mutations associated with increased viral fitness [62], while S235F (KSS = 0.79) is an Alpha-defining mutation. The ORF1ab position R5716C (KSS = 0.87), corresponding to NSP13 R392C in the helicase domain, distinguishes all post-BA.1 Omicron sublineages [63].

For HIV-1, the twelve top-ranking positions (KSS = 0.82–0.97) distributed across all three genes: seven in pol, four in gag, and one in env. The selection comprised three indels and nine substitutions, reflecting the indel constraint applied to a genome with extensive subtype-specific length polymorphisms.

In pol, KSS identified positions spanning four functional regions of the Gag-Pol polyprotein: transframe peptide (TFP), protease, reverse transcriptase (RT), and integrase. The highest-scoring pol position (KSS = 0.97; W34del, W34R, R36G) localizes to the transframe peptide (TFP/p6*), a region involved in Gag-Pol polyprotein processing and protease autoactivation [64], where subtype-specific deletions and substitutions distinguish multiple subtypes. Position 5 (KSS = 0.83; D5N, D5I) also maps to the TFP, in a region exhibiting subtype-specific variation [64]. In the protease, position 76 in pol (corresponding to protease standard position 20; KSS = 0.94; K76I) maps to a site where K20R/M/N are documented as PI-selected mutations [65]. In RT, K328 (KSS = 0.85; corresponding to RT position 173) maps to the palm subdomain [66], while position 358 (KSS = 0.87; RT position 203) achieved high discriminative capacity primarily through synonymous variation distinguishing subtypes, with non-synonymous E359L restricted to group O. A709S/N/K (KSS = 0.85; RT position 554) maps to the RNase H domain, within a region implicated in nucleoside analogue resistance [66, 67]. In the integrase domain, K851Q (KSS = 0.83; integrase position 136) localizes to the catalytic core domain near the DDE active site motif (D64, D116, E152) [68]. Position K136 has been experimentally implicated in viral DNA binding [69], and the K136Q polymorphism is associated with prior antiretroviral drug exposure [70], as catalogued in the HIV Mutation Browser [71].

In gag, KSS identified positions in two functionally distinct domains. In the matrix (MA/p17), E107I (KSS = 0.89) maps to helix 5 of the matrix domain, spatially close to E99 within the same helical segment in the trimeric structure. E99 is essential for Env incorporation into virions [72], and substitutions such as E107I may influence MA structural stability or membrane association. Position Q127del/V128del (KSS = 0.94) falls within an HLA-restricted cytotoxic T-lymphocyte (CTL) epitope (positions 124–132) where Q127 exhibits subtype-specific polymorphism reflecting population-level HLA-driven immune selection [73]. In the p6 domain, S465del (KSS = 0.95), the highest gag position, and E460A (KSS = 0.85) map to the functionally critical region immediately downstream of the PTAP late domain required for endosomal sorting complex required for transport (ESCRT)-mediated budding via Tsg101 recruitment [74], adjacent to the FRFG motif essential for Vpr incorporation into virions [75]. This region exhibits subtype-specific insertions and deletions, with PTAP motif duplications in subtype C conferring replication advantages [76, 77].

In env, the single top-ranking position V271I/M/R (KSS = 0.82) maps to the gp120 C2 conserved region [78], where the amino acid variant shows strong subtype association: V271I predominates in CRF01 AE, C, D, and multiple BC recombinants, V271M in A1, A6, and CRF63 02A6, while V271R is exclusive to group O (S5 Table).

For HCMV, gene-specific analysis (ten positions per gene) showed concordance with established genotyping systems. In UL55, the two highest-ranking positions (KSS = 0.86; N456del, H455del, D463del) localized to the hypervariable region flanking the furin cleavage site at codon 461, where deletions match genotype-defining criteria documented by Chou & Dennison [43]: group 2 carries N456del, while group 3 carries H455del and D463del. Additional top-ranking positions at codons 51 and 66 revealed novel discriminative N-terminal loci not included in the classical gB genotyping scheme [43]. In UL73, top positions (KSS = 0.57–0.79) mapped predominantly to the N-terminal domain (codons 4–79), matching genotyping criteria established by Pignatelli et al. [44]. Key polymorphisms included genotype-defining deletions at E18 and S33, with E18del characterizing genotype 4 subgroups and S33del distinguishing genotype 2, within a region clinically associated with symptomatic congenital infection [79].

In US28, the top positions (KSS = 0.47–0.80) distributed across functionally distinct domains: N-terminal extracellular positions (E18L, D19A) map to the chemokine-binding domain implicated in immune evasion [42, 80], while C-terminal intracellular position R329W (KSS = 0.66) maps to the signaling-relevant C-terminal tail, a region implicated in constitutive G-protein coupling [81]. US28 contained exclusively substitution-based top positions (0% indels), contrasting with UL55 (30% indels) and UL73 (30% indels), reflecting gene-specific patterns of selective pressure on this viral chemokine receptor. Three positions (KSS = 0.50–0.57) consisted entirely of synonymous mutations achieving high discriminative capacity with zero mutational impact, demonstrating the framework’s ability to identify molecular epidemiology markers independent of protein-level consequences [82, 83].

Across all three viruses, top-ranked positions showed strong concordance with literature-documented functional and strain-distinguishing loci: 12/12 SARS-CoV-2 positions correspond to established variant-defining or immune escape sites, 12/12 HIV-1 positions map to documented drug resistance, structural, or immune selection regions, and HCMV positions consistently matched established genotype-defining polymorphisms, with two additional UL55 positions (codons 51 and 66) representing novel discriminative loci not captured by classical genotyping schemes (S4–S6 Tables). These results were achieved without direct encoding of site-specific functional annotations. The HCMV analysis provided additional validation through substantially expanded datasets: 443 US28 sequences versus 55 in Arav-Boger [42] (8.1-fold increase), 399 UL55 sequences versus 14 in Chou [43] (28.5-fold increase), and 646 UL73 sequences versus 42 in Pignatelli [44] (15.4-fold increase). Original genotyping criteria, established with limited sequence availability, remained applicable to this expanded diversity.

### Limitations and future directions

The protein component depends on the completeness of UniProt annotations, meaning recently characterized proteins or under-represented viral families with sparse annotation histories may receive underestimated scores. This limitation is inherent to any annotation-dependent framework and implies that KSS analyses should be periodically re-evaluated as databases are updated. The mutational component provides biophysical granularity for amino acid substitutions, but assigns a uniform configurable score to insertions and deletions (default 1.0) regardless of structural context. This treatment cannot distinguish between indels with different biological consequences, for instance, small in-frame indels in flexible loop regions versus functionally critical deletions in antigenic domains.

Several methodological choices also constrain the framework. The discriminative component evaluates each position independently, capturing individual discriminative power but not epistatic or combinatorial interactions between positions. Default parameters (*k* = 9, *t* = 0.25, equal component weights) performed well across all tested systems, but the equal-weight scheme and protein scoring category weights (S1 Table) were set based on domain knowledge rather than formal optimization, and their sensitivity to alternative configurations has not been systematically assessed. Results also depend on the choice of reference genome, as positional coordinates and mutation calls are defined relative to a single reference per viral system; alternative references could shift specific position rankings, though overall scoring patterns should remain stable for well-characterized mutations. The current implementation requires users to provide pre-extracted coding sequences rather than automatically extracting them from whole-genome data. Automating this step would require integrating codon-aware alignment tools, which remain challenging for highly variable sequences due to reduced accuracy with frequent indels and reading frame shifts, or high computational costs [40, 84].

Validation encompassed three classes of viruses representing complementary evolutionary dynamics (positive-sense RNA, retrovirus, double-stranded DNA), but generalization to all viral types remains to be demonstrated. Negative-sense RNA viruses and segmented genomes may exhibit different mutational patterns requiring parameter adaptation. Furthermore, functional validation relied on retrospective concordance with published literature rather than prospective prediction. However, for a prioritization framework, retrospective agreement with independently established functional sites is a common validation approach, and the consistent concordance observed across three evolutionarily distinct classes and families of viruses (12/12 for both SARS-CoV-2 and HIV-1) substantially reduces the likelihood of chance agreement. Formal quantification of false positive rates remains difficult in the absence of a complete ground truth, as high-scoring positions without current literature support may represent genuinely relevant sites not yet characterized. Prospective experimental validation would be needed to assess predictive value for such sites.

Several directions could extend this work. Incorporating multivariate position selection could capture combinatorial discriminative patterns that univariate analysis misses, and the bounded [0, 1] scoring provides a compatible foundation for such integration. Context-dependent indel scoring, informed by protein structural data or disorder predictions, would improve mutational impact assessment for insertions and deletions. KSS-selected *k*-mers could serve as biologically interpretable features for machine learning classifiers, complementing pattern recognition with functional context.Real-time surveillance pipelines could leverage KSS to flag sequences carrying high-scoring *k*-mer combinations for expedited characterization. Extension to metagenomic applications would enable variant detection in complex viral populations, though handling fragmentary and mixed-infection sequences requires additional development. Finally, formal sensitivity analysis of component weights across diverse classes or families of viruses would provide empirical guidance for application-specific configurations.

## Conclusion

This study presents KSS, a framework integrating discriminative power with biological relevance for viral *k*-mer prioritization. Through systematic validation across SARS-CoV-2, HIV-1, and HCMV, we demonstrated that KSS can identify functionally significant and strain-distinguishing genomic positions while providing standardized, interpretable values in the [0, 1] interval enabling direct cross-dataset comparison.

The three-component architecture addresses critical limitations of existing *k*-mer selection methods. The discriminative component achieved classification performance comparable to the best univariate feature selection methods (mean F1 = 0.880 vs. 0.877 for chi-squared) while additionally providing consistent score distributions across all classes of viruses. The MIYATA EVO matrix, optimized via genetic algorithm, improved biophysical property correlations by 28.4% over the original MIYATA matrix, enhancing assessment of viral mutations where functional consequences outweigh evolutionary frequency patterns. The protein component showed robust concordance with UniProt annotations (*ρ* = 0.900, *τ* = 0.777) while providing continuous granularity that enables finer prioritization than categorical systems. Functional validation confirmed that top-ranked positions correspond to established variant-defining and functionally characterized loci across all three classes of viruses (12/12 for SARS-CoV-2, 12/12 for HIV-1, and concordance with classical genotyping schemes for HCMV).Notably, these results were achieved without direct encoding of site-specific functional annotations, demonstrating the framework’s capacity to recover biologically relevant positions from general discriminative and biophysical principles alone.

By combining computational discrimination with biological context through standardized scoring, KSS provides a validated approach for *k*-mer prioritization with potential applications in variant surveillance, molecular epidemiology, and functional annotation. The modular design and interpretable score decomposition support applications ranging from routine strain classification to identification of emerging mutations with potential phenotypic impact. The framework’s computational efficiency, demonstrated by complete analysis of over 290,000 sequences across 11 genes in under 24 hours on a single workstation-class laptop (Intel Core i9-14900HX, 64 GB RAM), supports deployment in the context of large-scale molecular surveillance.

## Supporting information

**S1 Table. UniProt-based scoring system for functional importance and characterization level of proteins**. Ten annotation categories with differential weights reflecting their relative contribution to functional characterization. Categories include Gene Ontology terms (molecular function, biological processes, cellular component), protein existence evidence, structural features, protein-protein interactions, post-translational modifications, 3D structures, drug target information, and literature references. Gene Ontology functional terms receive maximum weights. For biological processes, Reactome pathway cross-references are additionally counted. Total maximum: 100 points.

**S2 Table. Top 20 positive discordance proteins (high KSS, low UniProt annotation score)**. Proteins with Δ > 0.30 where KSS substantially exceeds normalized UniProt score. Columns: UniProt accession, annotation score (raw and normalized), KSS score, discordance (Δ), protein name, organism, protein existence level, number of references, Gene Ontology annotations (MF, BP, CC), 3D structure availability, post-translational modifications, known interactions, and number of structural/functional features.

**S3 Table. Top 20 negative discordance proteins (high UniProt annotation score, low KSS)**. Proteins with Δ *<* −0.30 where normalized UniProt score substantially exceeds KSS. Same column structure as S2 Table.

**S4 Table. SARS-CoV-2 top-ranking positions: detailed mutations and indel statistics**. Top 12 genome-wide KSS positions (Sheet 1) expanded to one row per variant *k*-mer. Columns: rank, gene, gene position, KSS and component scores (discriminative, mutational, protein), reference *k*-mer, variant *k*-mer, amino acid changes with mutational impact scores, indel status, and variant/lineage distribution. The variant distribution column reports the percentage of sequences carrying each *k*-mer variant within each class (variant, subtype, or genotype); classes with prevalence below 1% are omitted for clarity and summarized as “*n* others (*<*1%)”. Sheet 2 provides per-gene indel statistics (ORF1ab, S, M, N, E): total positions analyzed, positions with indels, and indel percentage.

**S5 Table. HIV-1 top-ranking positions: detailed mutations and indel statistics**. Top 12 genome-wide KSS positions (Sheet 1) expanded to one row per variant *k*-mer with the same column structure as S4 Table. Sheet 2 provides per-gene indel statistics for gag, pol, and env.

**S6 Table. HCMV top-ranking positions: detailed mutations and indel statistics**. Top 10 per-gene KSS positions for UL55, UL73, and US28 (Sheet 1) expanded to one row per variant *k*-mer with the same column structure as S4 Table. Sheet 2 provides per-gene indel statistics.

## Acknowledgments

The authors thank Amine M. Remita for insightful discussions and constructive feedback. The authors acknowledge the use of publicly available sequence data from GenBank and the Los Alamos National Laboratory HIV Sequence Database.

## Author contributions

**Conceptualization:** DL, FC, ABD. **Data curation:** DL, FC. **Formal analysis:** DL. **Funding acquisition:** ABD. **Investigation:** DL. **Methodology:** DL, FC, ABD. **Project administration:** DL. **Resources:** DL. **Software:** DL. **Supervision:** ABD. **Validation:** DL, FC, ABD, HS, SG, IB. **Visualization:** DL. **Writing – original draft:** DL, FC. **Writing – review & editing:** ABD, HS, SG, IB.

## Ethics statement

This study exclusively used publicly available sequence data from GenBank and the Los Alamos National Laboratory HIV Sequence Database. No human subjects, animal experiments, or personally identifiable information were involved. No ethics approval was required.

## Data availability

All source code, the complete framework implementation, evaluation scripts for reproducing all reported results, the MIYATA EVO substitution matrix, and the datasets used in this study are publicly available under the MIT license in the KSS GitHub repository at https://github.com/bioinfoUQAM/KmerSignificanceScore. The repository includes a requirements file specifying all software dependencies and versions (Python, BioPython, NumPy, SciPy, scikit-learn, pandas).

